# Enhancing efficacy of the MEK inhibitor trametinib with paclitaxel in *KRAS*-mutated colorectal cancer

**DOI:** 10.1101/2024.09.05.611458

**Authors:** Susmita Ghosh, Fan Fan, Reid Powell, Yong Park, Clifford Stephan, E. Scott Kopetz, Lee M. Ellis, Rajat Bhattacharya

**Affiliations:** Department of Surgical Oncology, The University of Texas MD Anderson Cancer Center, Houston, Texas; Department of Translational Medical Sciences, Texas A&M University School of Medicine, Houston, Texas; Department of Gastrointestinal Medical Oncology, The University of Texas MD Anderson Cancer Center, Houston, Texas; Molecular and Cellular Oncology, The University of Texas MD Anderson Cancer Center, Houston, Texas

**Keywords:** colorectal cancer, KRAS, trametinib, paclitaxel, combination therapy

## Abstract

**Background:** *KRAS* is frequently mutated in the tumors of patients with metastatic colorectal cancer (mCRC) and thus represents a valid target for therapy. However, the strategies of targeting KRAS directly and targeting the downstream effector mitogen-activated protein kinase kinase (MEK) via monotherapies have shown limited efficacy. Thus, there is a strong need for novel, effective combination therapies to improve MEK-inhibitor efficacy in patients with *KRAS*-mutated mCRC.

**Objective:** Our objective was to identify novel drug combinations that enhance MEK-inhibitor efficacy in patients with *KRAS*-mutated mCRC.

**Design:** In this study, we performed unbiased high-throughput screening (HTS) to identify drugs that enhance the efficacy of MEK inhibitors *in vitro*, and we validated the efficacy of the drugs *in vivo*.

**Methods:** HTS was performed using 3-dimensional CRC spheroids. Trametinib, the anchor drug, was probed with 2 clinically ready libraries of 252 drugs to identify effective drug combinations. The effects of the drug combinations on CRC cell proliferation and apoptosis were further validated using cell growth assays, flow cytometry, and biochemical assays. Proteomic and immunostaining studies were performed to determine the effects of the drugs on molecular signaling and cell division. The effects of the drug combinations were examined *in vivo* using CRC patient-derived xenografts.

**Results:** HTS identified paclitaxel as being synergistic with trametinib. *In vitro* validation showed that, compared with monotherapies, this drug combination demonstrated strong inhibition of cell growth, reduced colony formation, and enhanced apoptosis in multiple *KRAS*-mutated CRC cell lines. Mechanistically, combining trametinib with paclitaxel led to alterations in signaling mediators that block cell cycle progression and increases in microtubule stability that resulted in significantly higher defects in the mitosis. Finally, the combination of trametinib with paclitaxel exhibited significant inhibition of tumor growth in several *KRAS*-mutant patient-derived xenograft mouse models.

**Conclusion:** Our data provide evidence supporting clinical trials of trametinib with paclitaxel as a novel therapeutic option for patients with *KRAS*-mutated, metastatic CRC.

## INTRODUCTION

Metastatic colorectal cancer (CRC) is the second leading cause of cancer-related deaths in the United States.^1^ *KRAS* proto-oncogene, GTPase (*KRAS*) is one of the most commonly mutated oncogenes in human cancers, including colorectal cancer. More than half of patients with metastatic CRC have mutations in the oncogenic driver gene *KRAS*.^2^ In CRC, *KRAS* mutations cluster in 4 hotspots (codons G12, G13, Q61, and A146), with the most frequent site of *KRAS* mutations being codon G12 (65% of mutations). Within this codon, *KRAS* G12D is the most frequent mutation (44% of mutations), followed by *KRAS* G12V (30% of mutations) and *KRAS* G12C (13% of mutations).^3^ Despite the need to develop therapeutic strategies to target *KRAS*, direct targeting of *KRAS* has been unsuccessful for nearly 3 decades. However, recent breakthroughs have led to the development of small molecules that selectively and irreversibly bind to *KRAS* G12C.^4^ On the basis of the demonstrated efficacy of *KRAS* G12C inhibitors, the Food and Drug Administration (FDA) has approved the use of the *KRAS* G12C inhibitors sotorasib and adagrasib in *KRAS* G12C–mutated non-small cell lung cancer.^5, 6^ Inhibitors targeting other specific mutations of *KRAS* including agents targeting G12D^7^ and G12S^8^ and pan-*KRAS* inhibitors targeting both wild-type and mutated *KRAS*^9, 10^ are under development.

Despite the success of *KRAS* G12C inhibitors in patients with NSCLC, they have not been effective in patients with metastatic CRC with *KRAS* G12C mutations^11, 12^ and disease recurrence when treated with *KRAS* G12C inhibitors.^13^ Activation of alternative/bypass pathways,^14^ adaptive feedback reactivation of RAS-mitogen-activated protein kinase signaling,^15^ and resistance due to selection of drug-resistant mutants have been suggested as mechanisms of resistance to these inhibitors.^16^ Combination therapies involving sotorasib with panitumumab,^17^ and adagrasib with cetuximab^18^ have demonstrated moderate improvements over single agent G12C inhibitors in patients with mCRC. This has led to a recent approval by the FDA of the combination of adagrasib and cetuximab for treatment of patients with mCRC with *KRAS*-G12C mutations.

An alternative approach to targeting *KRAS*-driven tumors is to indirectly inhibit downstream mediators of RAS, such as mitogen-activated protein kinase kinase (MEK).^19^ However, single-agent targeting of MEK has not been clinically effective, particularly in patients with mCRC, because the activation of alternative pathways eventually leads to resistance. While several combinations of phosphatidylinositol 3-kinase [PI3K]/protein kinase B [AKT]/mammalian target of rapamycin [mTOR] inhibitors,^20–22^ B-cell lymphoma-extra large [BCL-XL] inhibitors,^23^ cyclin-dependent kinase 4/6 [CDK4/6] inhibitors,^24, 25^ and autophagy inhibitors^26, 27^ with MEK inhibitors have entered clinical trials, some combinations have failed due to enhanced toxicity,^20–22^ and the results for others are awaited. Thus, there is a strong need to identify novel, effective combination therapies that improve the efficacy of MEK inhibitors in patients with RAS-mutated, metastatic CRC.

The purpose of our study was to identify novel drug combinations that enhance the efficacy of MEK inhibitors in patients with *KRAS*-mutated, metastatic CRC. We hypothesized that an effective combination therapy with a MEK inhibitor combined with either a chemotherapy agent or a targeted therapy agent can be identified using an unbiased high-throughput screening (HTS) approach. Our preclinical studies demonstrated that combining trametinib with paclitaxel is more efficacious than trametinib alone. Our results can serve as the basis for future clinical studies to determine the efficacy of this novel drug combination in patients with *KRAS*-mutated metastatic CRC.

## METHODS

### Culture of CRC cell lines

All *KRAS*-mutated human CRC cell lines used in this study were purchased from ATCC. The cell lines included HCT116 (RRID:CVCL_0291), LS174T (RRID:CVCL_1384), LoVo (RRID:CVCL_0399), SW620 (RRID:CVCL_0547), and SW480 (RRID:CVCL_0546). Cells were cultured in minimum essential medium with 10% fetal bovine serum (Atlanta Biologicals) and vitamins, nonessential amino acids, penicillin/streptomycin, sodium pyruvate, and L-glutamine (Thermo Fisher Scientific) as supplements. The cell lines used for the *in vitro* experiments were limited to 15 passages. Validation of the cell lines was done at The University of Texas MD Anderson Cancer Center Characterized Cell Line Core Facility before conducting the experiments and at least once every year during the study. Before plating for each *in vitro* experiment, a cellometer (Nexelom Bioscience) was used according to the manufacturer’s instructions to determine cell numbers and viability .

### 3D, high-throughput screening

All HTS studies were performed at the Center for Translational Cancer Research (RRID: SCR_022214) at the Institute of Biosciences and Technology, Houston, Texas, as described previously.^28^ Briefly, for the 3D, HTS screening, 4 *KRAS*-mutated CRC cell lines— HCT116 (seeding density, 190 cells/well), SW620 (3,000 cells/well), LS174T (1,500 cells/well), and LoVo (1,500 cells/well)—were seeded in ultra-low-attachment, round (U)-bottom plates (Corning, Cat# 3830). A multidrop dispenser (Thermo Fisher Scientific) was used to form spheroids. The cells were then incubated in a robotically integrated Cytomat 6,000-cell culture incubator (Thermo Fisher Scientific) at 37 °C with 5% CO_2_. Cells were allowed to form spheroids for 48 hours after plating.

For the drug-combination screening, we tested the drugs in 2 libraries consisting of 252 FDA-approved and phase I-III investigational drugs obtained from the National Cancer Institute and the Center for Translational Cancer Research at the Institute of Biosciences and Technology, respectively, both as single agents and in multipoint, pairwise combinations with trametinib. After 7 days of drug treatment, cell plates were leveled to 35 μL using a HydroSpeed plate aspirator (Tecan) and a CellTiter-Glo 3D cell viability assay (Promega, Cat No. G9681) according to the manufacturers’ instructions. Luminescence was measured on a Synergy Neo2plate reader (Agilent BioTek).

### Bliss synergy analysis

For high-throughput combinatorial screenings, varying ratios of the anchor and probe were tested. Using a support vector machine–based method, the data were fit to a 3D surface to provide additional rigor, and automated outlier detection was done as previously described.^29^ The Bliss independence model was then used to calculate the theoretical additivity surface.^30^ By comparing the empirically determined drug effect to the Bliss independence model, the interactions between each pair of drugs were characterized as antagonistic, additive, or synergistic. As a subjective cut-off, we used a volumetric difference (i.e., the sum of all pairwise interactions) of −1 or 1 to define antagonism or synergy, respectively.

### 3-(4,5-dimethylthiazol-2-yl)-2,5 diphenyltetrazolium bromide and colony formation assays

Initially, cell viability was measured using 3-(4,5-dimethylthiazol-2-yl)-2,5 diphenyltetrazolium bromide (MTT) reagent as described previously.^31^ Briefly, CRC cells were plated at a seeding density of 1,000-3,000 cells/well in 96-well, flat-bottom plates. After 24 hours, cells were treated with trametinib (5 nM), paclitaxel (5 nM), or both drugs (Selleck Chemicals). Dimethyl sulfoxide was used as the control. After 48 hours of treatment with the drugs, cell viability was measured by adding MTT reagent and performing colorimetric measurements using a plate reader.

For the colony formation assays, CRC cells were plated at a seeding density of 1,000-3,000 cells/well in 12-well dishes. Cells were treated with different doses of trametinib or paclitaxel alone for 7 days, and the half-maximal inhibitory concentrations for each drug/dose combination were determined. For the combination studies, both trametinib and paclitaxel were used at concentrations lower than their half-maximal inhibitory concentrations, and the cells were cultured for 7 days. The drugs were used as single agents at the same concentrations for comparison. Cells treated with dimethyl sulfoxide were used as controls. Methylene blue solution (0.05%) was used to stain the surviving colonies, which were imaged. Then, sodium dodecyl sulfate (1%) was used to extract the cell-bound dye, and the intensity of the colored solutions was measured at 600 nm.

### Western blot analyses

HCT116 (0.1 × 10^6^ cells/well) and SW620 cells (0.3 × 10^6^ cells/well) were seeded in 6-well plates. The next day, the cells were treated with trametinib (5 nM), paclitaxel (10 nM), or both for 24 or 48 hours. Cell lysates were prepared in a radioimmunoprecipitation assay buffer with protease and phosphatase inhibitors as described previously.^32^ The extracted proteins were separated using sodium dodecyl sulfate polyacrylamide gel electrophoresis following a standard protocol and transferred to Immobilon polyvinylidene membranes (EMD Millipore). For 1 hour, the membranes were subjected to blocking with 5% milk in tris(hydroxymethyl)aminomethane (Tris) buffered saline with 0.1% Tween 20 (TBST); then, they were incubated overnight at 4 °C with primary antibodies diluted in 3% bovine serum albumin in TBST. After being washed in TBST (3 times), the membranes were reincubated with horseradish peroxidase–labeled secondary antibodies for 1 hour. The membranes were again washed in TBST (3 times) and exposed to autoradiography films. Signals were detected using chemiluminescence. Antibodies for cleaved poly(ADP-ribose) polymerase (PARP; Cat# 9541; RRID:AB_331426), cyclin-dependent kinase inhibitor 1B (P27-Kip1; Cat# 2552; RRID:AB_10693314), retinoblastoma protein (pRB; Cat# 8516; RRID:AB_11178658), forkhead box protein M1 (FOXM1; Cat# 5436; RRID: AB_10692483), and phospho-histone H2AX (pH2AX; Cat# 9718; RRID:AB_2118009) were from Cell Signaling Technologies, and those for acetylated-tubulin (Cat# 23950; RRID:AB_628409) and α-tubulin (Cat# 32293; RRID:AB_628412) were from Santa Cruz Biotechnology. All antibodies were used according to the manufacturers’ instructions.

### Reverse-phase protein array

HCT116 (0.1 × 10^6^ cells/well) and SW620 cells (0.3 × 10^6^ cells/well) were seeded in 6-well plates. The next day, the cells were treated with trametinib, paclitaxel, or both and were allowed to grow for 48 hours. The preparation of cell lysates was done as described above. Reverse-phase protein array (RPPA) analyses were performed at MD Anderson’s Functional Proteomics (RPPA) Core Facility as previously described.^32^

### Immunofluorescence staining

For immunofluorescence staining, HCT116 cells (0.1 × 10^6^ cells/well) were seeded in 6-well plates containing 4 coverslips. Cells were cultured for 24 hours and were then treated with trametinib (10 nM) and paclitaxel (5 nM) alone or in combination for 48 hours. The coverslips were further processed for immunofluorescence staining. The cells were washed in phosphate-buffered saline, fixed in methanol, rehydrated in TBST, and incubated with either anti-acetylated tubulin or α-tubulin antibodies diluted in TBST + 3% bovine serum albumin. Following washing with TBST, the coverslips were incubated with goat anti-mouse Alexa-594 secondary antibodies (Cat# A-11032; RRID:AB_ 2534091). The nuclei were stained and visualized using Hoechst 33342 (Thermo Fisher Scientific). Images were obtained using a fluorescence microscope (Olympus BX71) using either 40X or 60× objective lenses. For presentation, acetylated tubulin and α-tubulin were colored green and DNA was colored red using Adobe Photoshop version 2004.

### *In vivo* studies using patient-derived xenografts

The mice used for the *in vivo* studies were acquired from the Department of Experimental Radiation Oncology at MD Anderson. Three CRC patient-derived xenografts (PDXs)—PDX C1117 (*KRAS* G12D), PDX C1138 (*KRAS* G13D), and PDX B8239 (*KRAS* G12C)—were used in this study. The PDXs were initially grown subcutaneously in male Nod/SCID/gamma mice as described previously.^24, 28^ When the tumors reached approximately 1 cm in diameter, the mice were euthanized and the tumors were harvested. The tumors were then cut into small (approximately 2-mm) pieces using a sterile scalpel and were implanted subcutaneously into the flanks of anesthetized 4–6-week-old nude mice. When the tumors reached approximately 100-200 mm^3^, the mice were randomly distributed into 4 treatment groups: vehicle, trametinib (0.2 mg/kg, 5 days/week, given orally), paclitaxel (10 mg/kg, 2 times/week, given intraperitoneally), or trametinib plus paclitaxel [dosage, schedule, and administration routes as above] for about 3-4 weeks. For PDX C1117 and PDX B8239, there were 10 mice per treatment group. However, for PDX C1138, the take rate and growth kinetics of the tumors were different; therefore, mice were randomized to 5 per treatment group to ensure similar tumor sizes at the start of the experiment.

Trametinib (Selleck Chemicals) was prepared in a suspension of 0.5% H-methyl cellulose plus 0.5% Tween 80, and paclitaxel (Division of Pharmacy, MD Anderson) was prepared in 0.9% saline solution. Digital calipers were used to measure tumor sizes, and mice were weighed twice a week by a blind observer. All animal experiments were performed under a protocol approved by the Institutional Animal Care and Use Committee at MD Anderson. The PDXs were obtained from a repository at MD Anderson through a collaboration with Dr. E. Scott Kopetz.

### Statistical analyses

GraphPad Prism 9 and Microsoft Excel were used to create graphical representations of the study results. For the *in vitro* assays, all quantitative values represented at least 3 replicates. For the *in vivo* assays, 8-10 tumors were measured for each treatment group. Two-tailed Student *t*-tests were used to compare groups. The results were expressed as means plus or minus the standard errors of the means (SEMs). *P* < 0.05 was considered significant.

## RESULTS

### High-throughput screening identified paclitaxel was synergistic with trametinib in multiple *KRAS*-mutated CRC spheroids

Unbiased HTS was performed using 252 FDA-approved and phase I-III investigational drugs from 2 drug libraries; the National Cancer Institute Oncology Set V and a custom clinical drug set available from the Center for Translational Cancer Research at the Institute of Biosciences and Technology, respectively. Trametinib was used as the base compound, and the drugs in the libraries were used either as single agents or in combination with trametinib. We used a 3D, spheroid-based HTS assay that we had previously developed^28^ to identify agents that enhanced the efficacy of trametinib, either synergistically or additively, in *KRAS*-mutated CRC cell lines (Figure 1A). Based on its effects on CRC cell growth, which was in excess of the Bliss score for synergy, the antimitotic, microtubule-targeting agent paclitaxel was identified as one of the drugs that synergistically enhanced the efficacy of trametinib at clinically relevant concentrations (Figure 1B). Our initial observations and further validation studies demonstrated drug synergy between paclitaxel and trametinib in 4 CRC cell lines with *KRAS* mutations: SW620 (*KRAS* G12V), HCT116 (*KRAS* G13D), LoVo (*KRAS* G13D), and LS174T (*KRAS* G12D). Collectively, these initial data provided a strong rationale to further investigate the effects of paclitaxel in combination with trametinib in additional model systems.

**Figure 1:**
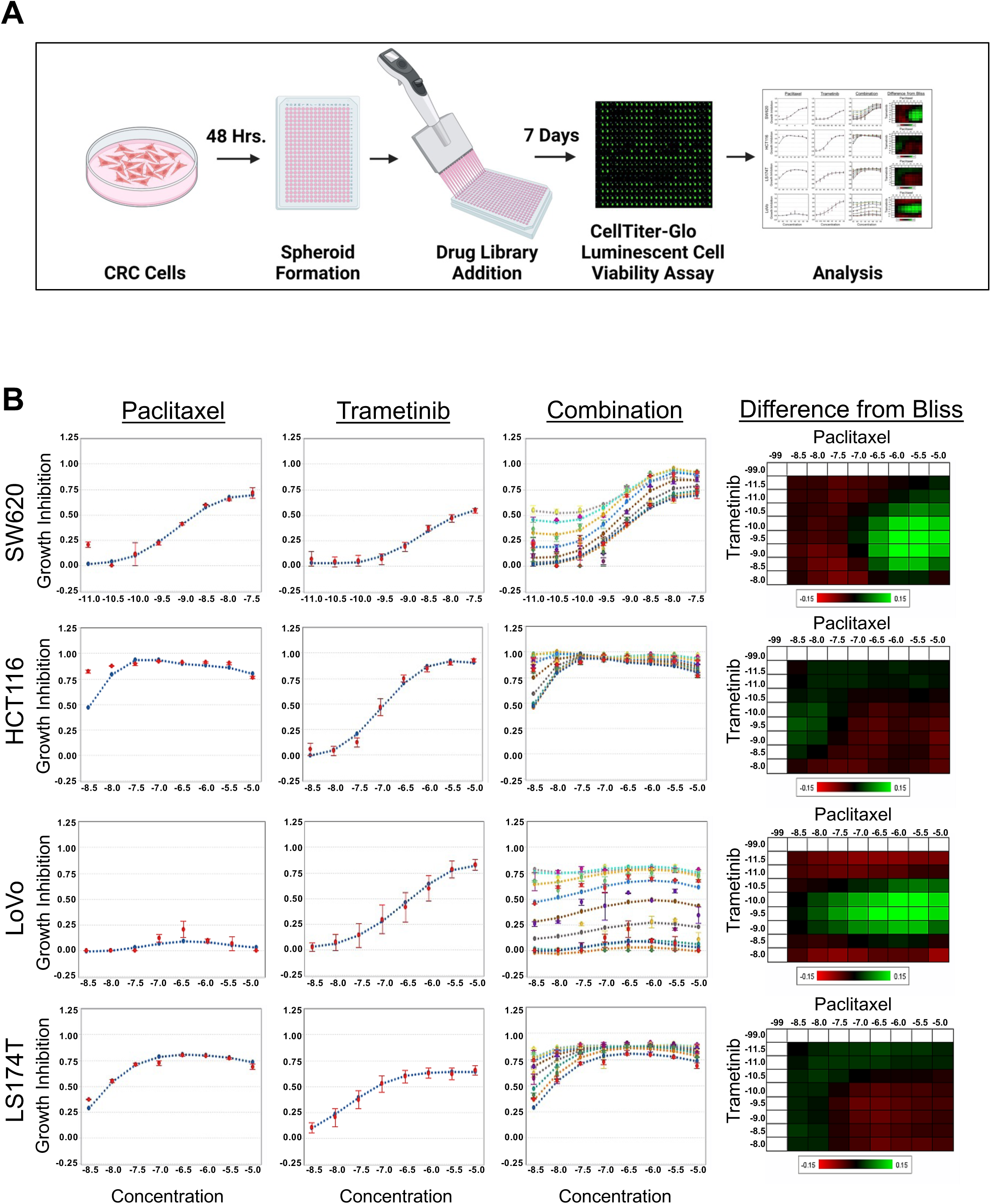
High-throughput screening indicated that paclitaxel was synergistic with trametinib in multiple *KRAS*-mutated CRC spheroids. **(A)** Schematic illustration of HTS using 3D spheroids to identify drugs synergistic with trametinib. Images were created using BioRender. **(B)** HTS using SW620, HCT116, LS174T, and LoVo 3D spheroids. Panels from left to right: 3D spheroid growth curves for the probe drug paclitaxel as a single agent, for the anchor drug trametinib as a single agent, and for paclitaxel plus trametinib at the indicated concentrations based on CellTiter-Glo assays and heat maps showing drug synergy as measured by the volumetric difference from that determined using the Bliss independent model. Green areas represent a difference from Bliss > 0, indicating synergy. Synergy was observed at multiple drug doses (green areas). The drug concentrations are shown on a log scale. Data are presented as the mean ± the SD. 3D, 3-dimensional; CRC, colorectal cancer; Hrs., hours; HTS, high-throughput screening; SD, standard deviation.

### Paclitaxel enhanced the efficacy of trametinib in multiple *KRAS*-mutated CRC cell lines

The combination of trametinib and paclitaxel was further evaluated for its effects on colony-forming ability in a panel of *KRAS*-mutated CRC cells. The results from the clonogenic assay demonstrated that the combination of trametinib and paclitaxel was more effective than either drug as a single agent and that it significantly suppressed cell proliferation, as measured by the reduction in the number of proliferating colonies in the SW620, HCT116, SW480, and LS174T cell lines (Figure 2A). Measurement of the extracted methylene blue in each well also supported that the drug combination was more effective in inhibiting CRC cell growth as compared to single agents alone.

**Figure 2:**
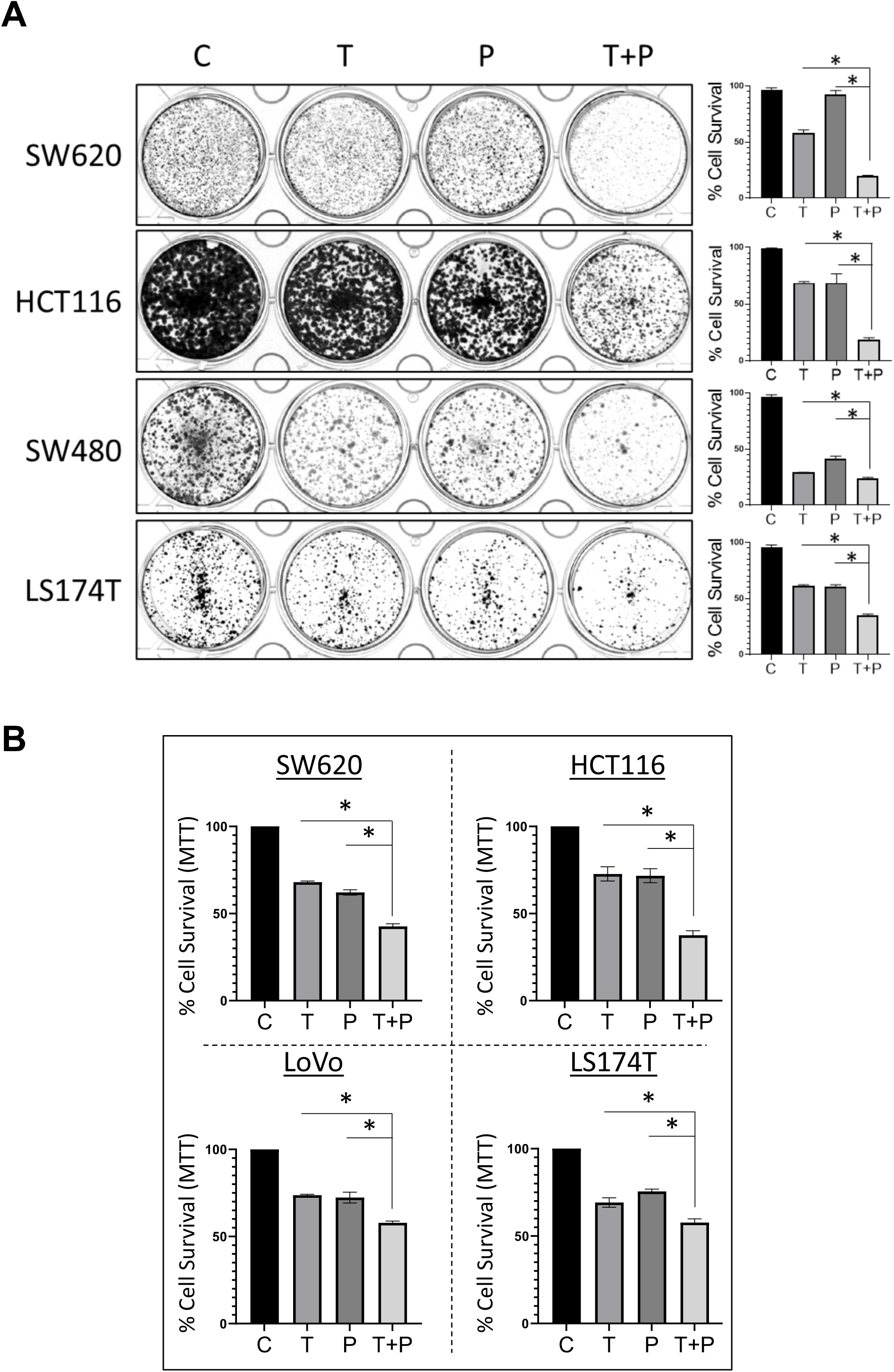
The combination of trametinib and paclitaxel was synergistic in multiple *KRAS*-mutated CRC cell lines. **(A)** Colony formation assays evaluating the synergy of trametinib and paclitaxel in multiple *KRAS*-mutated CRC cell lines. The left panel shows representative images of methylene blue–stained colonies for each cell line examined. The graphs on the right show the fraction of surviving cells based on optical density measurements of methylene blue absorbed by CRC cells. All data are presented as means ± SEMs. \**P* < 0.05 compared to cells treated with the control or single agents (Student *t*-test). **(B)** MTT assays validating the synergistic reduction of cell survival in the indicated *KRAS*-mutated CRC cell lines. Graphs showing survival of cells by treatment type. All data are presented as means ± SEMs. \**P* < 0.05 compared to trametinib alone (Student *t*-test). C, untreated control; CRC, colorectal cancer; MTT, 3-(4,5-dimethylthiazol-2-yl)-2,5 diphenyltetrazolium bromide; P, paclitaxel; SEM, standard error of the mean; T, trametinib; T+P, trametinib plus paclitaxel.

To further validate the effectiveness of the combination of trametinib and paclitaxel on cell proliferation, we performed MTT assays in multiple *KRAS*-mutated CRC cell lines. The combination inhibited cell proliferation more than either drug alone or dimethyl sulfoxide control in the SW620, HCT116, LS174T, and LoVo cell lines (Figure 2B).

### The combination of trametinib and paclitaxel enhanced cell death in *KRAS*-mutated CRC cell lines

Next, in the SW620 and HCT116 cell lines, we used flow cytometry and Western blot analysis to examine the effect of combining trametinib and paclitaxel on cell death by the induction of apoptosis. Flow cytometric staining demonstrated the presence of significantly higher levels of total apoptotic cells (cells positive for Annexin V and for both Annexin V and propidium iodide) in sets treated with the drug combination as compared with untreated sets or those treated with a single agent (Figure 3A). Western blot analysis (Figure 3B) demonstrated an increase in cleaved PARP in CRC cells treated with the drug combination as compared with untreated or trametinib-treated cells. However, PARP cleavage was not significantly increased in the presence of the drug combination as compared with paclitaxel only. Overall, these studies demonstrated an increase in CRC cell death in the presence of the trametinib/paclitaxel combination, but they suggest that apoptosis is not the only mode of cell death induced by the combination drug treatment.

**Figure 3:**
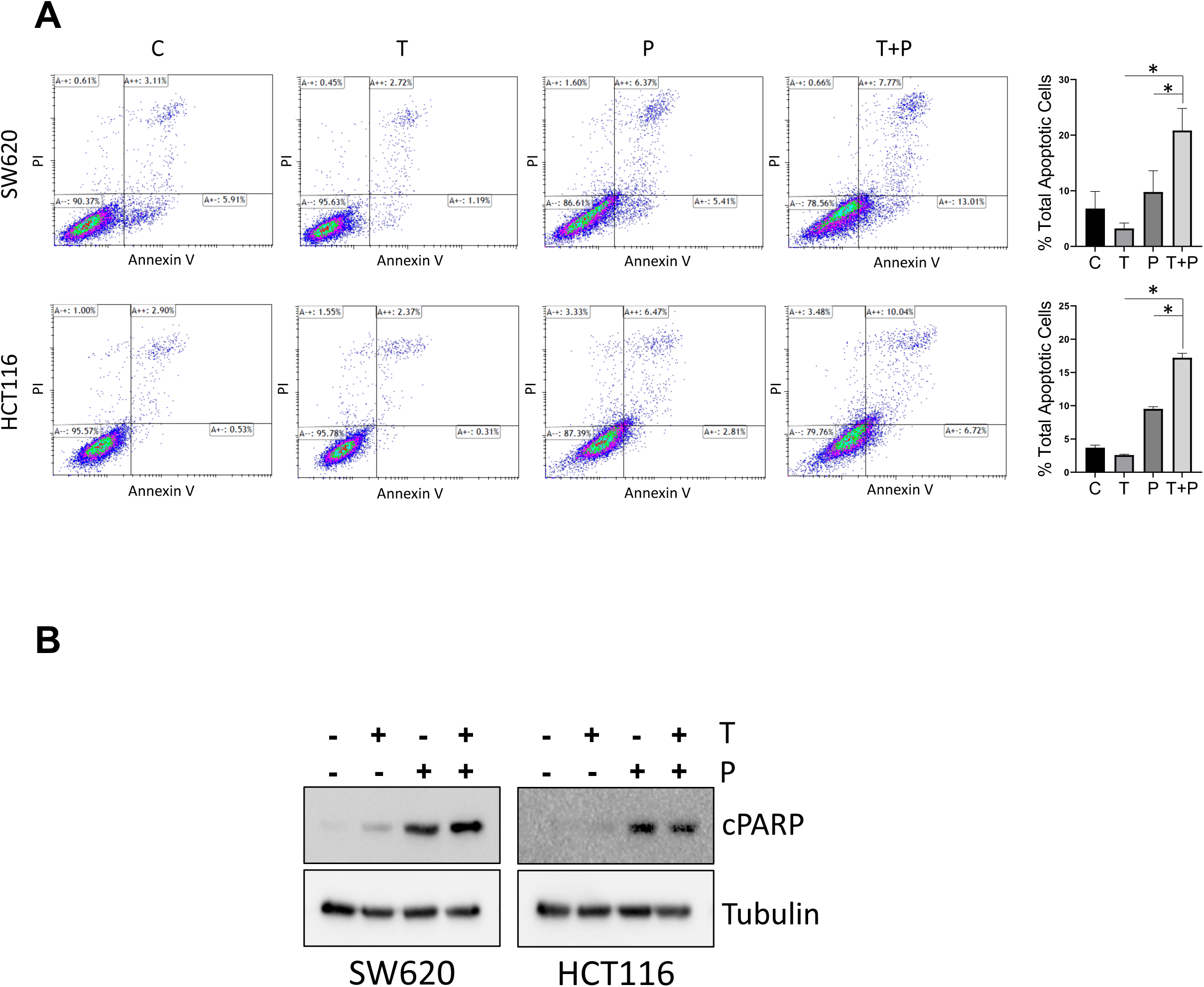
The combination of trametinib and paclitaxel enhanced cell death in *KRAS*-mutated CRC cell lines. **(A)** Flow cytometric analysis showing increased cell death in *KRAS*-mutated CRC cell lines treated with trametinib plus paclitaxel. SW620 and HCT116 cells were treated with trametinib (10 nM), paclitaxel (5 or 10 nM), and their combination for 72 hours. Data for each group are given in box plots. The graphs on the right indicate the total percentage of apoptotic cells (positive for Annexin V or for both Annexin V and propidium iodide) in each group. All data are presented as means ± SEMs. \**P* < 0.05 compared to treatment with the control or single agents (Student *t*-test). **(B)** Representative Western blots showing the protein levels of markers of apoptosis (cleaved PARP) in SW620 and HCT116 cells treated with trametinib, paclitaxel, or both for 48 hours. Tubulin was the loading control. C, untreated control; cPARP, cleaved poly (ADP-ribose) polymerase; CRC, colorectal cancer; P, paclitaxel; SEM, standard error of the mean; T, trametinib; T+P, trametinib plus paclitaxel.

### The combination of trametinib and paclitaxel inhibited cell-cycle progression and induced DNA damage in *KRAS*-mutated CRC cell lines

To determine the effect of the drug combination on cellular signaling that leads to a reduction in CRC cell proliferation, we performed a RPPA analysis using protein lysates from the SW620 and HCT116 cell lines. The RPPA analyses demonstrated significantly higher levels of P27-Kip1 and lower levels of cyclin B, CDK1, pRB, and FOXM1 in the cells treated with the drug combination as compared with the untreated cells or those treated with single agents (Figure 4A). Western blot analyses confirmed significantly higher levels of P27-Kip1 and lower levels of pRB and FOXM1 in the combination-treated cells compared with the cells treated with monotherapies (Figure 4B). A significant increase in the induction of DNA damage in the presence of the drug combination was demonstrated by increases in the pH2AX levels in both the RPPA and Western blot analyses (Figure 4A and 4B). Together, these results demonstrate significant alterations in the markers of cell-cycle progression in *KRAS*-mutated CRC cells treated with the drug combination compared with single agents.

**Figure 4:**
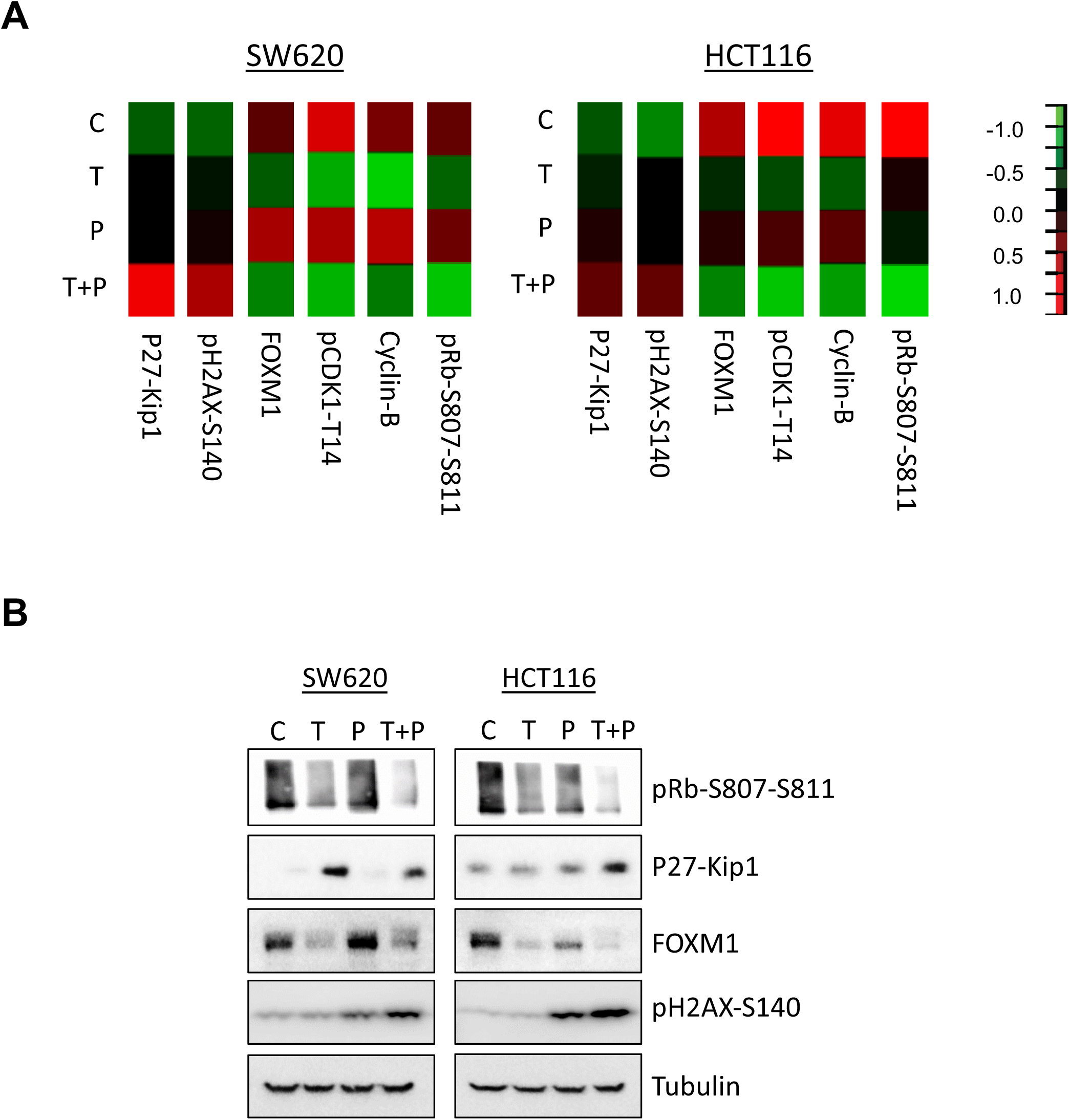
The combination of trametinib and paclitaxel inhibited cell-cycle progression and induced DNA damage in *KRAS*-mutated CRC cell lines. **(A)** Heat maps from the RPPA analysis demonstrating the inhibition of cell-cycle progression and DNA damage from treatment with trametinib plus paclitaxel in *KRAS*-mutated CRC cell lines. Only selected proteins are shown. SW620 and HCT116 cells treated with 5 nM trametinib, 10 nM paclitaxel, or both for 48 hours. (B) Representative Western blots validating the decrease in the markers of cell-cycle progression and DNA damage following treatment with the combination of trametinib and paclitaxel in SW620 and HCT116 cells. C, untreated control; CDK1, cyclin-dependent kinase 1; cPARP, cleaved poly (ADP-ribose) polymerase; CRC, colorectal cancer; FOXM1, forkhead box protein M1; P, paclitaxel; P27-Kip1, cyclin-dependent kinase inhibitor 1B; pH2AX, phospho-histone H2AX; pRB, phospho-retinoblastoma; RPPA, reverse-phase protein array; T, trametinib; T+P, trametinib plus paclitaxel.

### Trametinib enhanced the effects of paclitaxel in *KRAS*-mutated CRC cell lines

Paclitaxel enhances the stability of microtubules and induces mitotic defects in cancer cells.^33, 34^ However, its effects can be reduced by the increased function or overexpression of ABCB/ABCG transporter proteins, which results in the increased clearance of paclitaxel from cancer cells.^35, 36^ Our studies and studies from other investigators have indicated that trametinib can negatively affect the function of these transporter proteins^28, 37^ and thus potentially increase the intracellular concentration of paclitaxel. We hypothesized that the presence of trametinib increases the concentration of paclitaxel inside CRC cells, increasing microtubule stability and enhancing mitotic defects. Western blotting of cell lysates and immunostaining of fixed cells demonstrated a significant increase in the levels of acetylated tubulin, a validated marker for microtubule stability, in CRC cells treated with the drug combination as compared with single-agent treatments (Figure 5A and 5B). We next calculated the number of mitotic cells with normal bipolar, defective bipolar, monopolar, and multipolar mitotic spindles in the presence of paclitaxel, trametinib, or both. The immunofluorescence results demonstrated a significantly higher number of cells with defective bipolar, monopolar, and multipolar mitotic spindles in HCT116 cells treated with the combination of trametinib and paclitaxel as compared with either drug alone (Figure 5C). Compared with treatment with single agents and control, treatment with the drug combination resulted in about 2- to 3-fold and 9-fold increases, respectively, in the total number of defective mitotic cells.

**Figure 5:**
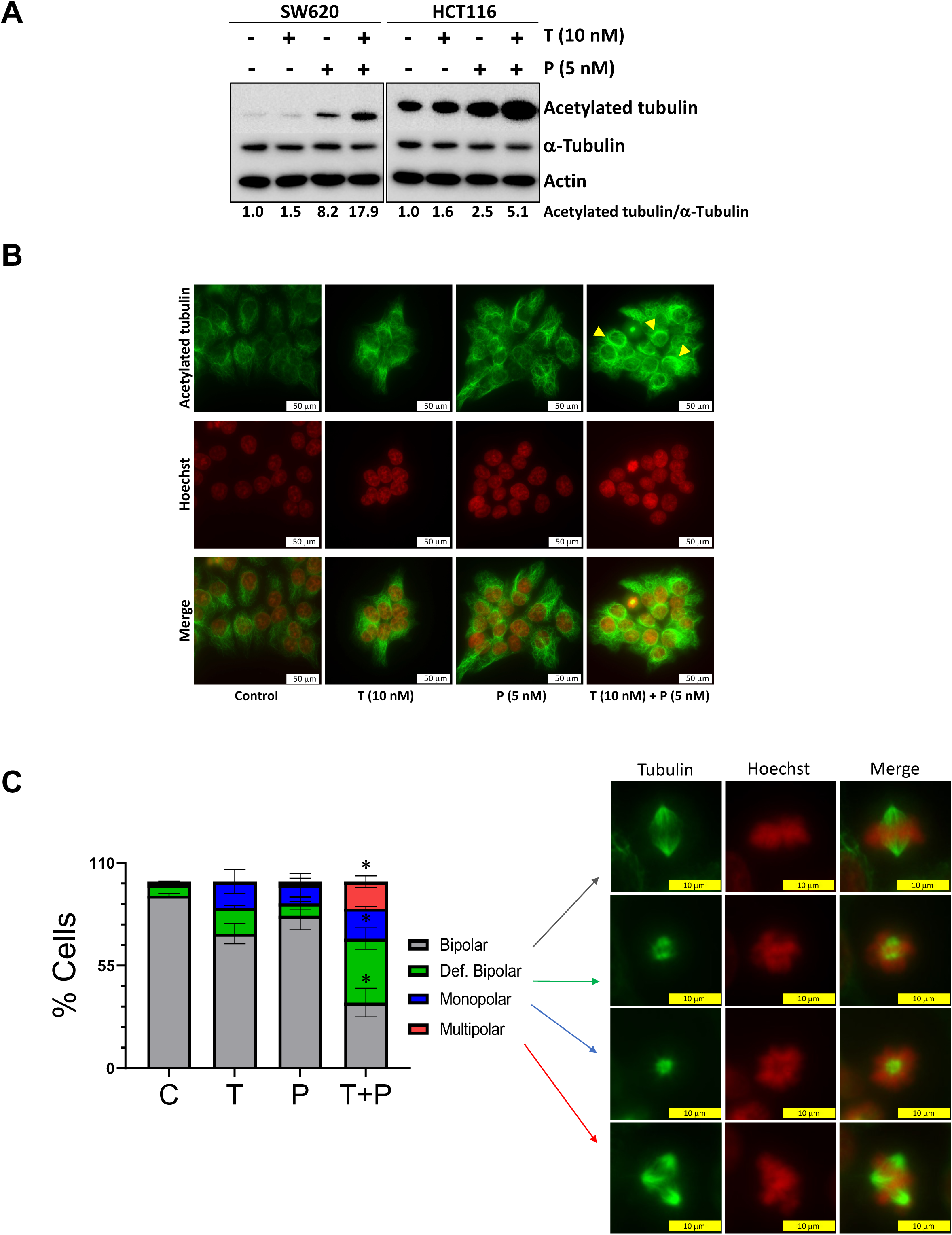
The combination of trametinib and paclitaxel increased mitotic defects in *KRAS*-mutated CRC cells. **(A)** Western blot to measure the levels of acetylated tubulin and α-tubulin in lysates of CRC cells treated with trametinib, paclitaxel, or both for 48 hours. Actin was the loading control. The numbers below the blots denote the levels of acetylated tubulin relative to α-tubulin for each sample. **(B)** Fluorescent images of fixed CRC cells treated with trametinib, paclitaxel, or both for 48 hours. Green indicates acetylated tubulin, and red indicates nuclei. The arrows point to microtubule bundles. Scale bar = 50 μm. **(C)** The graph in the left panel shows the increased percentage of defective mitotic cells (with bipolar, defective bipolar, monopolar, and multipolar spindles) in HCT116 cells treated with trametinib plus paclitaxel compared with those treated with trametinib, paclitaxel, or the control. Representative images of various mitotic cells are shown in the right panels. Microtubules are shown in green and DNA is shown in red. Scale bar = 10 μm. Only HCT116 cells were used for the immunostaining studies, as the fixation of the mitotic SW620 cells was difficult and unreliable. C, untreated control; CRC, colorectal cancer; def, defective; P, paclitaxel; T, trametinib; T+P, trametinib plus paclitaxel.

### Trametinib in combination with paclitaxel inhibited tumor growth in KRAS-mutated, patient-derived xenografts

To validate the antitumor effects observed in our *in vitro* studies, we treated mice bearing PDXs with 3 different *KRAS* mutations with trametinib, paclitaxel, or both drugs. The mice were treated with animal-equivalent doses that were calculated using published guidelines for the doses approved for human subjects.^38^ In the C1117 (*KRAS* G12D) and C1138 (*KRAS* G13D) PDX models, the tumor volumes were significantly decreased in the mice treated with the combination of trametinib and paclitaxel compared with those treated with the control or with single agents (Figure 6A and 6B). In the B8239 (*KRAS* G12C) PDX model, the average tumor volume was significantly decreased (by approximately 50%) in the mice treated with the combination of trametinib and paclitaxel compared with those treated with the control or single-agent paclitaxel (Figure 6C). However, no significant reduction in tumor volume was observed when mice treated with the combination therapy were compared with those treated with trametinib. To determine if the drug combination was well-tolerated, the body weights of the mice were assessed throughout the study. Combination treatment with trametinib and paclitaxel was not associated with a significant decrease in the mice’s weights (Figure 6A, 6B, and 6C).

**Figure 6:**
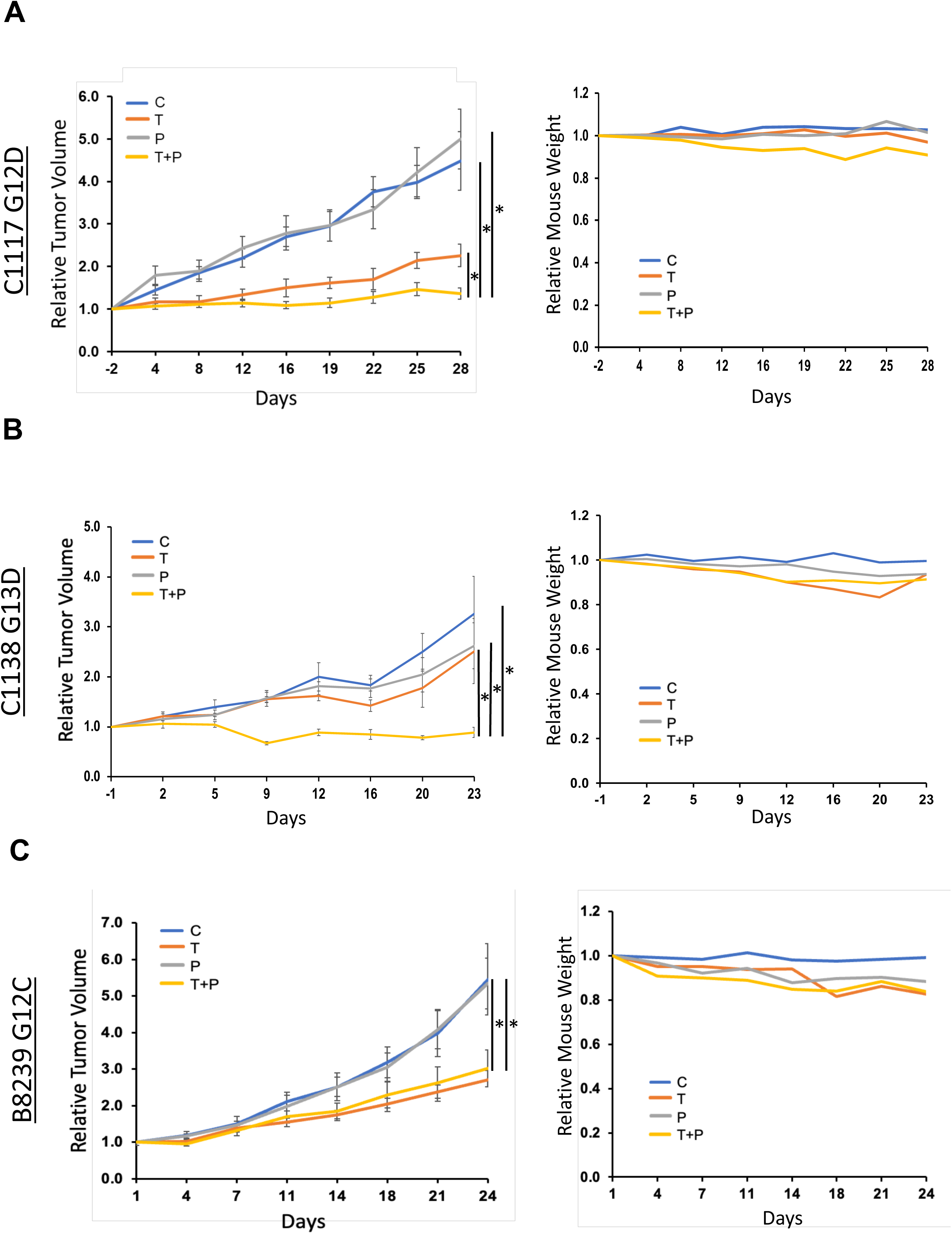
Trametinib in combination with paclitaxel inhibited tumor growth in multiple *KRAS*-mutated, patient-derived xenografts. *KRAS*-mutated PDXs (C1117, C1138, and B8239) were grown subcutaneously in mice and treated with the control, trametinib (0.2 mg/kg, oral, 5 days/week), paclitaxel (20 mg/kg, IP, twice/week), or trametinib plus paclitaxel **(A, C, E)** Left panels; Plots demonstrating relative tumor growth. Tumor volumes were measured on the indicated days (x-axes). Day 1 indicates the start of treatment. Data are presented as means ± SEMs. \**P* < 0.05 compared to treatment with the control and the single agents (Student *t*-test). Right panels: Plots of the relative weights of the mice measured on the days indicated in the x-axes. C, untreated control; P, paclitaxel; PDX, patient-derived xenograft; SEM, standard error of the mean; T, trametinib; T+P, trametinib plus paclitaxel.

## DISCUSSION

In this study, we present evidence that combining paclitaxel with trametinib enhanced the efficacy of trametinib in preclinical models of *KRAS*-mutated CRC. The combination not only led to increased cytotoxicity in *KRAS*-mutated CRC cells *in vitro*, but also suppressed tumor growth in *KRAS*-mutated CRC PDX murine models with no evidence of significant drug-related toxicity.

*KRAS* has been targeted for decades, with efforts culminating in the recent development of inhibitors that specifically target *KRAS* G12C.^39, 40^ Novel agents targeting other *KRAS* and pan-*KRAS* mutations have also been developed and are now in various stages of preclinical or clinical assessment for efficacy in various cancers, including CRC.^7–10^ While *KRAS* G12C inhibitors have demonstrated efficacy in non-small cell lung cancer, they have shown limited efficacy as monotherapies in patients with *KRAS*-mutated CRC. Studies have shown that patients treated with these inhibitors develop drug resistance over time through multiple mechanisms,^39, 40^ including reactivation of the RAS-mitogen-activated protein kinase signaling pathway.^15^ Initial reports on combining *KRAS* G12C inhibitors with anti–epidermal-growth-factor-receptor antibodies have reported modest improvements in progression-free survival in patients with *KRAS* G12C–mutated, metastatic CRC.^18, 41, 42^

Because directly targeting *KRAS* in CRC with single agents has been ineffective, targeting *KRAS*-mutated CRCs with MEK inhibitors as single agents has also been investigated. However, these MEK inhibitors were ineffective as single agents^43^ due to various mechanisms of resistance,^44^ highlighting the need for therapeutic approaches that could potentially augment the sensitivity of CRC tumors to MEK inhibitors as a component of a multidrug regimen. We performed unbiased HTS screening to identify potential combinatorial therapeutic regimens using the MEK inhibitor trametinib (FDA-approved for treating patients with melanoma) as the backbone to treat *KRAS*-mutated CRC tumors. Because 3D cultures represent the tumor microenvironment better than 2D cultures,^45^ we used 3D CRC spheroids in the HTS studies to identify clinically relevant drug combinations. Candidate drugs were drawn from 2 different library sets. We identified paclitaxel as being synergistic with trametinib in *KRAS*-mutated CRC spheroids. We then validated our HTS results using different *in vitro* approaches in multiple *KRAS*-mutated CRC cell lines. Both clonogenic assays to measure long-term exposure and MTT assays to measure short-term exposure to the drug combination resulted in significantly reduced colony formation and cell proliferation in CRC cell lines with different *KRAS* mutations, suggesting increased cytotoxicity from the drug combination. To evaluate the cause of the increased cytotoxicity, we examined apoptotic proteins and cell-cycle markers. Using flow cytometry, we observed an increase in the total number of apoptotic cells in the combination group compared with the single-agent and control groups.

The drug combination also affected cell-cycle regulation. It is known that, during the G1 phase of the cell cycle, phosphorylation of the retinoblastoma protein (RB) leads to the activation of E2F proteins and the expression of E2F target genes. This cluster of genes encodes cell-cycle regulators, such as cyclin A, cyclin E, and CDK1. During the G2 phase of the cell cycle, CDK2/cyclin A and CDK1/cyclin B complexes serially phosphorylate FOXM1, leading to the activation of FOXM1 target genes. This cluster of genes encodes cell-cycle regulators such as cyclin B and centromere protein-F that is required for the execution of mitosis.^46^ In this study, RPPA and Western blot analyses showed that, compared to cells treated with monotherapy, cells treated with the combination of paclitaxel and trametinib showed decreased pRB, CDK1, cyclin B, and FOXM1 levels. Cells treated with the paclitaxel/trametinib combination also showed increased levels of P27-kip1, an inhibitor of cell-cycle progression. These results indicate that, *in vitro*, the drug combination effectively inhibits cell-cycle progression.

Mitotic catastrophe, which can be induced by DNA damage or mitosis errors, is defined as a cellular mechanism that stops cancerous cells from proliferating and as a mode of cell death occurring after improper progression of the cell cycle. Paclitaxel is an antimicrotubule agent that causes mitotic catastrophe by binding to microtubules and altering their dynamics,^34, 47^ resulting in hyperstabilized and dysfunctional microtubules *in vitro*.^47, 48^ This results in mitotic spindle dysfunction, chromosome missegretation, and finally mitotic catastrophe, leading to cell cycle arrest or cell death.^34, 49^ Our study showed that paclitaxel treatment enhanced the stability of microtubules in CRC cells compared to control cells; this was established by the presence of acetylated tubulin, a known marker for stable microtubules. Interestingly, trametinib treatment also consistently increased the stability of microtubules, albeit to a lesser extent, in treated as compared with control cells. However, in all the CRC cell lines examined, paclitaxel-induced microtubule stability was very strongly enhanced in cells treated with both paclitaxel and trametinib compared with untreated cells or cells treated with paclitaxel alone. These novel findings demonstrated a significant enhancement of paclitaxel activity when the drug was combined with trametinib. Additionally, as predicted, there was a significant increase in the number of CRC cells exhibiting various mitotic defects among cells treated with trametinib plus paclitaxel compared with untreated cells or those treated with a single agent, and this finding points to a clear mechanism underlying the synergy of this drug combination.

Paclitaxel is widely used in the treatment of various cancers, but it is not approved for the treatment of patients with CRC most likely due to high level of expression of the multi drug resistance protein (MDR1). The overexpression or increased activity of ATP-binding cassette (ABC) transporters, most notably ABCB1, is one of the foremost drivers of multidrug resistance in cancer cells.^50^ Expression of these transporter proteins in CRC cells^51^ can thus likely remove multiple cytotoxic agents including paclitaxel from cancer cells and result in reduced intracellular concentrations and efficacy of these drugs and increased drug resistance.^35, 36^ Our studies and other previous studies have shown that trametinib can inhibit the function of ABCB1 and thus enhance the intracellular concentrations of multiple cytotoxic drugs.^28, 37^ Based on the enhanced activity of paclitaxel in the presence of trametinib, we suggest that trametinib inhibits the function of transporter proteins, leading to an increase in intracellular paclitaxel. Increased paclitaxel levels result in enhanced mitotic defects that, in combination with the inhibitory effects of paclitaxel on various cell cycle regulators, augment the cytotoxic effects of trametinib, leading to drug synergy.

In summary, we performed unbiased, 3D HTS studies and found that paclitaxel is synergistic with the MEK inhibitor trametinib in *KRAS*-mutated CRC. This drug combination led to significantly decreased cell proliferation, increased apoptosis, and inhibited cell-cycle progression *in vitro*. It also significantly suppressed tumor growth *in vivo* in different *KRAS*-mutated PDXs, and mice tolerated the regimen well. Therefore, our preclinical studies show that this drug combination is effective, and our results support its further study in clinical trials to determine its efficacy in patients with *KRAS*-mutated metastatic CRC.

## STATEMENTS AND DECLARATIONS

### Ethical considerations

All animal studies were performed following approval of the study protocol by the Institutional Animal Care and Use Committee (IACUC) at the University of Texas MD Anderson Cancer Center at Houston.

### Consent to participate

Not applicable.

### Consent for publication

Not applicable.

### Data availability

The datasets generated and analyzed during this study has been included in the manuscript.

## Acknowledgement

We thank Laura L. Russell, scientific editor, Research Medical Library, for editing this article.

